# Increased Activity of Tyrosine Hydroxylase Leads to Elevated Amphetamine Response and Markers of Oxidative Stress in Transgenic Mice

**DOI:** 10.1101/188318

**Authors:** Laura M. Vecchio, M. Kristel Bermejo, Amy Dunn, Marija Milenkovic, Nikhil Urs, Amy Ramsey, Gary W. Miller, Ali Salahpour

## Abstract

In Parkinson’s disease, noradrenergic cells of the locus coeruleus and dopamine cells within the nigrostriatal pathway undergo profound degeneration. Tyrosine hydroxylase (TH) is the rate-limiting enzyme in the production of all catecholamines, including dopamine and noradrenaline, and is selectively expressed in the cells that produce these neurotransmitters. *In vitro* studies have previously shown that the TH-synthetic system can contribute to the formation of reactive oxygen species. In addition, animal models of dopamine mishandling demonstrated that free dopamine is neurotoxic. To examine how increased TH activity might influence catecholamine systems *in vivo*, we generated TH-overexpressing mice (TH-HI) with six total copies of the *TH* murine gene. A commensurate increase in *TH* mRNA produced a threefold increase in both total TH protein and phosphorylated TH levels. We found an increased rate of dopamine synthesis in both young and adult mice, reflected by the accumulation of L-DOPA following NSD-1015 administration, as well as elevated dopamine tissue content in young mice and an increased presence of dopamine metabolites at both ages. Adult mice show no difference in baseline locomotor behaviour compared to wildtype littermates, but a have potentiated response to amphetamine. In addition to elevated dopamine turnover in the striatum, TH-HI mice show reduced levels of glutathione and increased levels of cysteinylated catechols. These results indicate that a heightened level of active TH can produce oxidative stress, and may represent a source of toxicity that is specific to catecholamine cells, which are most vulnerable to degeneration in Parkinson’s disease.

## Introduction

The neuropathology of Parkinson’s disease is marked by the degeneration of dopamine and noradrenaline neurons as well as the accumulation of alpha-synuclein protein aggregates. By the time its characteristic motor impairments have manifested, approximately 80% of cells within the nigrostriatal dopaminergic pathway have been lost. Importantly, dopaminergic loss is preceded by comparable degeneration of noradrenergic cells in the locus coeruleus, which likely underlies many of the early non-motor symptoms of Parkinson’s disease (Braak et al., 2003; Fahn, 2003). While oxidative stress is recognized to play a key role in pathogenesis of Parkinson’s disease and in the mechanisms of neurodegeneration, why dopaminergic and noradrenergic cells are most profoundly affected is still the subject of investigation (Zhang et al., 2000; Miyazaki and Asanuma, 2008).

A source of oxidative stress unique to cells that produce catecholamines, such as dopamine and noradrenaline, is the small amounts of reactive oxygen species (ROS) produced as by-products of catecholamine synthesis — a reaction that is mediated by the rate-limiting enzyme tyrosine hydroxylase (TH) (Adams et al., 1997; Haavik, 1997; Haavik et al., 1997; Haavik and Toska, 1998). Under normal conditions, ROS produced by the TH biosynthetic system can be neutralized by glutathione and superoxide dismutase (Perry and Yong, 1986; Spina and Cohen, 1989). However, an increase in TH activity may accelerate ROS production. Past studies have suggested that in synucleinopathic disorders, increased enzymatic activity may come as a result of diminished regulation placed on TH (Perez and Hastings, 2004; Alerte et al., 2008; Farrell et al., 2014). Both *in vitro* and *in vivo* studies have demonstrated that alpha-synuclein has the capacity to negatively regulate the activity of TH (Perez et al., 2002; Perez and Hastings, 2004; Peng et al., 2005; Park et al., 2007; Alerte et al., 2008; Liu et al., 2008; Farrell et al., 2014). Indeed, mouse models with mutated or aggregated alpha-synuclein have been shown to have increased levels of TH activity, as well as behavioural impairment attributed to catecholamine dysregulation (Alerte et al., 2008; Farrell et al., 2014). In *Drosophilia* and *C. elegans*, dysfunctional alpha-synuclein was shown to increase levels of phosphorylated TH and elevate dopamine levels, and ultimately resulted in oxidative stress and neurodegeneration (Park et al., 2007; Locke et al., 2008).

Importantly, the accumulation of dopamine in the cytosol has itself been identified as a source of oxidative stress and neurotoxicity: left unsequestered, cytosolic dopamine can auto-oxidize to produce quinones, superoxide and H2O2, and/or cysteinyl adducts (Hastings et al., 1996; Chen et al., 2008; Miyazaki and Asanuma, 2008; Stansley and Yamamoto, 2013). Transgenic mouse models have demonstrated that increased uptake of dopamine from the extracellular space can lead to increased dopamine turnover and cell loss in the nigrostriatal system, as can an impairment in the ability to sequester intracellular dopamine into vesicles (Caudle et al., 2007; Masoud et al., 2015). Like these models of dopamine mishandling, it is possible that an accumulation of cytosolic dopamine might also result from poorly regulated dopamine synthesis.

We hypothesize that an increase in the enzymatic activity of TH has the capacity to produce disruptions in catecholamine homeostasis *in vivo* and as a consequence, the potential to contribute to conditions of oxidative stress. To evaluate the potential of heightened catecholamine synthesis to contribute to oxidative stress in brain regions most affected by Parkinson’s disease, we developed a novel line of transgenic mice overexpressing functional TH. While previous studies have attempted to develop TH overexpressing mice (Kaneda et al., 1991), our model is the first to successfully achieve functional overexpression of TH and increased synthesis in endogenous catecholaminergic regions.

## Materials and Methods

All experiments used age-and sex-matched wildtype and transgenic mice maintained on a C57Bl/6J genetic background. Experiments were performed in junvenile mice at 4 weeks and adult mice at approximately 12 weeks of age (range: 10-20 weeks). Mice were housed on a 12 hour light-dark cycle, with food and water provided *ad libitum*. Procedures conformed to the recommendations of the Canadian Council on Animal Care, with protocols reviewed and approved by the University of Toronto Faculty of Medicine and Pharmacy Animal Care Committee.

### Generation Of Tyrosine-Hydroxylase Overexpressing Mice

Mice overexpressing TH were created by bacterial artificial chromosome (BAC) transgenesis using an artificial chromosome that contains the murine TH locus as well as approximately 90 kb of genomic DNA upstream and downstream to the gene (RP23-350E13, Genome Sciences). The genomic DNA carried one other gene sequence (*Tspαn32*), a transcription factor (Ascl-2), and three other predicted genes and predicted pseudogenes (R74862, Gm39117, Gm7290), and a portion of the sequence for gene *Cd81*. While these sequences were noted, they were not believed to bear consequences for our transgenic model: *Tspαn32* is specific to haematopoietic tissues and not expressed in the brain, Ascl-2 requires dimerization with other proteins to function, and the predicted genes have no known functions. In addition to the murine genomic DNA, the RP23-350E13 vector sequence contains a gene encoding chloramphenicol resistance. Bacterial colonies were grown on a chloramphenicol-spiked lysogeny broth (LB) agar plate, and single colonies were randomly selected for growth in liquid LB media (12.5 μg/mL chloramphenicol). The colonies were verified by PCR-based genotyping using primers that amplify a sequence overlapping the BAC vector and the *TH* gene (forward, *5’-gaaggctctcaagggcatc;* reverse, *5’-acaggtcaggctctcaggtc*).

The BAC DNA was isolated using a NucleoBond BAC 100 preparation kit and resuspended in injection buffer (0.03 mM spermine/0.07 mM spermidine). The DNA preparation was diluted to 2 ng/μl in injection buffer before *in vitro* pronuclear microinjection of fertilized eggs, which were then implanted into pseudo-pregnant C57Bl/6 surrogate mothers. Pronuclear microinjections were performed at Emory University transgenic facility. Transgenic positive pups were identified by PCR-based genotyping (*5’-aggagctgactgggttgaag;* reverse, *5’-tcgtggcctgttgtgagtag*); again, primer sequences overlapped the vector and gene sequences, thereby differentiating transgenic TH from native TH. Of the F1 generation, a mouse determined by qPCR to possess six copies of genomic *TH* was used as a founder for a transgenic high copy line, TH-HI. Transgenic mice were maintained on a C57 background.

### Determination Of Gene Copy Number And Total TH mRNA Levels

#### gene copy number

Genomic DNA was isolated from tail biopsy after Proteinase K digestion and isopropanol precipitation (Burkhart et al., 2002; Ballester et al., 2004), and cleaned using a chloroform-ammonium acetate wash method. Optical density was measured at 260/280 nm, and dilutions of both 20 ng/ul and 5 ng/ul were prepared from each DNA sample.

Quantitative PCR (qPCR) was used to determine total allelic copy number of TH, inclusive of both native alleles as well as transgenic copies (Ballester et al., 2004; Shepherd et al., 2009; D’Haene et al., 2010). Briefly, amplicons of the *TH* genomic sequence were generated using designed primers (forward: 5’-caccagtcctgagtttcctatt, reverse: 5’-ctggatcacactccaccatatc) and measured relative to those of a common reference gene that encodes the transferrin receptor (*Tfrc*; forward: 5’-cagtcatcagggttgcctaata, reverse: 5’-atcacaacctcaccatgtaact). Relative quantification of the gene targets was obtained at both dilutions using the GoTaq® qPCR Master Mix (Promega, Fitchburg, WI, USA) and the Applied Biosystems 7500 Real-Time PCR System. The PCR protocol used for all reactions was as follows: 2 minutes at 50 °C, 10 minutes at 95 °C, 40 repeats of 95 °C for 15 seconds, and 1 minute at 60 °C. Analysis of gene copy number was based upon the 2^-∆∆CT^ method, as previously described (Livak and Schmittgen, 2001; Pfaffl, 2001; Schmittgen and Livak, 2008).

#### mRNA

Reverse-transcriptase quantitative PCR was also used to determine total levels of mRNA. Following cervical dislocation, brains were removed and frozen in 2-methylbutane over dry ice, with each placed in an individual 15 mL RNA-free tube. Whole brains were stored at −80 C and subsequently used for RNA extraction. Midbrain regions were dissected from a 1 mm coronal section homogenized in cold Tri-Reagent (BioShop), and processed as previously described (Hu et al., 2009). Optical density readings at 260/280 nm were used to determine the quality and concentration of RNA. For reverse transcriptase PCR, 300 ng of RNA was used to obtain complementary DNA using SuperScript III Reverse Transcriptase (Invitrogen) according to manufacturer’s instructions. As before, quantification of mRNA levels was obtained using the GoTaq® qPCR Master Mix on the Applied Biosystems 7500 Real-Time PCR System. To ensure that the level of *TH* mRNA was normalized to the volume of dopamine tissue within the dissected area, *TH* mRNA levels (forward primer: *5′-cgggcttctctgaccaggcg*; reverse primer: *5′-tggggaattggctcaccctgct*) were presented as a ratio of *DAT* mRNA levels (forward primer: *5′-ggcctgggcctcaacgacac*; reverse primer: *5′-ggtgcagcacaccacgtccaa*). Relative quantification of both gene targets was made against the reference gene, phosphoglycerate kinase 1 (PGK1; forward primer: *5’-ggcctttcgacctcacggtgt*; reverse primer: *5′-gtccaccctcatcacgacccg*), using the ∆ ∆∆Ct method.

### Western Blot

Western blots were performed to quantify TH and DAT protein expression, using the sodium-potassium pump (Na^+^/K^+^-ATPase) as a loading control. The midbrain and striatum were dissected from freshly harvested brains and mechanically homogenized in RIPA buffer [50 mM Tris·HCl, pH 7.4; 150 mM NaCl; 1% Nonidet P-40; 0.5% sodium deoxycholate; 0.1% sodium dodecyl sulfate (SDS)] with protease inhibitors (working concentrations: pepstatin A, 5 μg/mL; leupeptin, 10 μg/mL; aprotinin, 1.5 μg/mL; benzamidine, 0.1 μg/mL; PMSF, 0.1 mM). Samples were spun at 21,130 × g (15,000 rpm) for fifteen minutes after homogenization, and protein concentrations were determined using a bicinchoninic acid (BCA) assay (Pierce, 23225). Samples were prepared to the desired loading concentration (striatum, 25 μg; midbrain, 60 μg) in a solution containing 1*x* SDS sample buffer (4 *x* solution: 3.4 g Tris Base, 8.0 g SDS, 40 mL glycerol in 1 L dH2O, pH-ed to 6.8 with concentrated HCl), 0.05% β-mercaptoethanol (BME), and RIPA buffer. Preparations were then heated to 55 °C for ten minutes, separated by 10% SDS/PAGE, and transferred to polyvinylidene difluoride (PVDF) membranes. Membranes were blocked in Li-COR buffer for one hour (room temperature) before incubating overnight (4 °C) with rabbit anti-Na^+^/K^+^-ATPase (1:2000; Cell Signaling, 3010S) and rat anti-DAT (1:750; Chemicon, MAB369) in the same buffer. After washing, protein bands were revealed by incubating with the fluorescence-labelled secondary antibodies donkey anti-rat IRdye 800CW and goat anti-rabbit AlexaFluor AF680 (1:5000; Rockland, 612-732-120 and Invitrogen, A21076, respectively). Membranes were then stripped and re-probed with rabbit anti-TH (1:3000 dilution; Millipore, AB152) in 5% non-fat milk, which was again revealed with goat anti-rabbit AlexaFluor AF680.

Using similar methods, TH protein levels in both central and peripheral noradrenergic regions were examined. Tissue from the TH-rich locus coeruleus and the adrenal medulla was extracted, homogenized in RIPA buffer and spun at 21,130 × g prior to sample preparation. Samples were again run on 10% SDS/PAGE gels (locus coerleus, 60 μg; adrenal medulla, 20 μg), transferred to PVDF membranes, and probed for rabbit anti-TH. Mouse anti-tubulin (Developmental Studies Hybridoma Bank, E7) and mouse anti-GAPDH as loading controls for the locus coeruleus and adrenal medulla (respectively), and revealed with donkey anti-mouse IRdye 800CW (1:5000 dilutions; Rockland, 610-731-002).

In a separate set of experiments, striatal samples were speedily dissected and homogenized in RIPA containing phosphatase inhibitors (final concentrations: sodium pyrophosphate, 2.5 mM; β-glycerophosphate, 1.0 mM; sodium fluoride, 50 mM; sodium orthovandadate, 1.0 mM) in addition to the protease inhibitors listed above. Membranes were incubated overnight (4°C) in solutions containing rabbit anti-phosphoTH-Ser19 (1:2000; Phospho Solutions, p1580-19), rabbit anti-phosphoTH-Ser31 (1:1000; Phospho Solutions, p1580-31) or rabbit anti-phosphoTH-Ser40 (1:2000; Phospho Solutions, p1580-40). Proteins were revealed using goat anti-rabbit AF680 (1:5000) before being stripped, and re-probed using mouse anti-TH (1:2500; Sigma, T2928) and mouse anti-GAPDH in 5% non-fat milk.

Westerns were also performed on 4-week old and/or 12-week old mice to assess levels of the vesicular monoamine transporter (VMAT2) using pre-cast gels and MOPS (3-(N-morpholino) propanesulfonic acid) buffers, with rabbit anti-VMAT2 antibody (1:10,000) generated by Dr. Gary Miller’s laboratory (Caudle et al., 2007; Lohr et al., 2014). Protein was revealed with goat anti-rabbit 800 (1:5000; Rockland, 611-131-002) by incubating for one hour at room temperature, and membranes were re-probed for TH and GAPDH as described above.

Densitometric analyses were conducted using Image Processing and Analysis in Java (ImageJ) software (http://rsb.info.nih.gov/ij/index.html).

### Immunohistochemistry

Immunohistochemistry was performed on coronal sections 40 μm cut from the brains of adult mice. In sections containing the midbrain and striatum, TH and DAT were immunostained using methods previously described (Vecchio et al., 2014); TH and the noradrenaline transporter (NET) were immunostained in the locus coeruleus. *Primary antibodies*: rabbit anti-TH (1:2000), rat anti-DAT (1:200), and anti-NET (1:500; MAb Technologies, NET05-2). *Secondary antibodies*: anti-rabbit AF680 (4000), anti-rat IRdye 800 (1:2000), and anti-mouse IRdye 800CW (1:2000).

## Tissue Content

### Dopamine and Metabolites

Total tissue content levels of dopamine and its metabolites, 3,4-dihydroxyphenylacetic acid (DOPAC) and homovanillic acid (HVA), were quantified in the striatum using high-performance liquid chromatography (HPLC). For all tissue content analyses, striata were dissected and flash frozen in liquid nitrogen, each hemisphere in a separate eppendorf tube. Only one side (the left or right striatum) was used for tissue content experiments, unless otherwise noted.

Tissue content analyses were conducted using an AgAgCl-electrochemical detector (LC-4C Amperometric Detector; BASi) set at an oxidizing potential of + 0.75 V, and a Hypersil Gold C18 column (150 × 3 mm, particle size of 5 μm; Thermo Scientific). The mobile phase contained 24 mM Na2HPO4, 3.6 mM 1-octanesulfonic acid, 30 mM citric acid, and 0.14 mM EDTA in 19% methanol and was adjusted to pH 4.7 using concentrated NaOH. After allowing the column to equilibrate with the mobile phase, the separation of pure standards was verified (100 ng/mL in HClO4): dopamine; DOPAC; HVA; serotonin; 5-hydroxyindoleacetic acid (5-HIAA), a metabolite of serotonin; and 2,3-dihydroxybenzoic acid (DHBA), an internal standard (all chemical standards from Sigma, purity > 98%). If separation of the compounds was not satisfactory, minor adjustments were made to the pH using HCl or NaOH, or to the methanol content; when adjustments to the mobile phase were required, the column would again be allowed equilibrate. Once separation of standards was achieved, a calibration curve was constructed at eight dilutions, each of which contained dopamine, DOPAC, HVA and DHBA (5, 10, 25, 50, 75, 100, 250, 500 ng/mL). Following successful calibration (1 ≥ R^2^ ≥ 0.995 for each standard), unilateral striata were weighed while frozen and homogenized in 0.1 M HClO4 spiked with 100 ng/mL DHBA (40 μl per mg of tissue), then centrifuged 10,000 x *g* for 10 minutes at 4 °C. The supernatant was carefully removed and filtered through a 0.22 μm membrane (Ultrafree-MC GVWP filters; Millipore, UFC30GV00), again by centrifugation at 10,000 x *g* (2 minutes, 4°C). The tissue content of dopamine, DOPAC and HVA was quantified against their respective calibration curves, and normalized to DHBA. The tissue content was quantified in two age cohorts: young mice, 4-5 weeks of age, and adults, 10-20 weeks of age. Two batches of samples were run (each containing mice of both genotypes at both ages), and therefore normalized to levels of 4-week wildtype mice within their respective cohort.

### L-DOPA

In a separate set of experiments, striatal tissue content of L-3,4-dihydroxyphenylalanine (L-DOPA) was measured to confirm an increase in functional TH in transgenic animals. A single IP injection of NSD-1015 (100 mg/kg), a compound known to inhibit the activity of dopamine decarboxylase, was administered to prevent the rapid conversation of L-DOPA to dopamine (Jones et al., 1998b). The extent of L-DOPA accumulation in striatal samples is accepted as a reliable measure of total TH enzymatic activity *in vivo* (Ferris et al., 2014), and therefore considered a measure of the rate of catecholamine synthesis. After 40 minutes, mice were sacrificed by cervical dislocation and the striatum were dissected and flash frozen in liquid nitrogen. Striatum were weighed and homogenized in 40 μl per mg sample of 0.1 M HClO4 spiked with both DHBA and 100 ng/mL sodium metabisulphate (Moron et al., 2000; Caudle et al., 2007), and prepared as described in the preceding paragraph. L-DOPA and stores of dopamine, DOPAC, and HVA were separated by HPLC and measured by electrochemical detection, using the methods described in the preceding section. The mobile phase was composed of 5.99 mg monobasic sodium phosphate, 200 μM EDTA, 0.684 g sodium chloride, 60 mg octyl sodium sulfate, and 95 mL of methanol in 1 L HPLC-grade water (pH 2.43). Analytes were quantified against standard curves, as previously described, with the addition of an L-DOPA standard (Sigma, D9628).

### Cysteinyl - DOPAC and Cysteinyl-L-DOPA

Cysteinyl analytes were measured as previously reported (Caudle et al., 2007). Briefly, bilateral striata were sonicated for 10 seconds in 0.1M perchloric acid containing 347μM sodium bisulfite and 134μM EDTA (1:6, w/v), then centrifuged at 10,000 x *g* for 10 minutes. Here, supernatant was filtered at 0.22 μm at 10,000 x *g* for 12 minutes. An ESA 5600A CoulArray detection system, equipped with an ESA Model 584 pump and an ESA 542 refrigerated autosampler was used for analysis (ESA, Bedford, MA). Separations were performed at 24.5°C using an XTerra MS 4.6 x 150mm 5μM C18 column. The mobile phase consisted of 0.65 mM 1-octanesulfonic acid sodium, 75 mM NaH2 PO4, and 8% acetonitrile (pH 2.20). Samples were eluted isocratically at 0.8 mL/minute and detected using a 6210 electrochemical cell (ESA) equipped with 5020 guard cell. Again, analytes were identified by the matching criteria of retention time and sensor ratio measures to known standards.

### Glutathione Assay

Decreased levels of glutathione have been correlated to a rise in oxidative stress, as well an increase in protein carbonyls (Dalle-Donne et al., 2003; Aldini et al., 2006; Chen et al., 2008). As an indication of oxidative stress, glutathione levels in the striatum and the cortex (a negative control) were assessed using a glutathione assay kit (Cayman, 703002). Brains were removed and quickly rinsed in iced cold PBS to remove blood on the exterior of the brain; striatal and cortical samples were then dissected and flash frozen in liquid nitrogen. Dissected tissue was weighed while frozen, and homogenized in ice-cold buffer (0.4 M 2(N-morpholino) ethanesulphonic acid, MES; pH 6-7, containing 1mM EDTA; 10 μl per mg of tissue). Samples were homogenized and spun at 10,000 x *g* for 15 minutes at 4°C, and the supernatant removed. The supernatant was deproteinated using metaphosphoric acid (MPA) and 4 M triethanolamine (TEAM) solution, as per kit instructions. The assay was performed as directed, and the absorbance measured at 5 minute intervals for 30 minutes (405 and 414 nm). Glutathione concentrations were determined using the kinetic method. As intraperitoneal injections of reserpine are known to induce oxidative stress and reduce glutathione in dopaminergic cells (Spina and Cohen, 1989), a subgroup of wildtype mice were injected with 5 mg/kg of reserpine (injection volume: 0.2 mL/g) two hours prior to dissection, as a positive control of oxidative stress (Spina and Cohen, 1989).

### Locomotor Activity and Amphetamine Challenge

Locomotor activity was measured in automated locomotor activity monitors. In these experiments, total distance covered and stereotypic behaviours was collected in 5 minute bins over the course of the test period. Mice were placed into the activity monitor chamber and measured at ‘baseline’ conditions for 60 minutes. Following baseline recording, mice were injected with a dose of saline, 2 mg/kg or 3 mg/kg amphetamine and recorded for an additional 90 minutes (IP, injection volume of 0.1 mL/g) (Salahpour et al., 2008; Vecchio et al., 2014).

### Statistical Analyses

All statistical analyses were performed for each group by Student’s t-test, one-way or two-way ANOVA. *Post-hoc* analyses were performed using the Bonferroni test (all pairwise comparisons). Significance is reported where *P* < 0.05.

## Results

### Production of TH-Overexpressing Mice Using BAC Transgenesis

Purified DNA containing the murine *TH* gene was microinjected into pseudopregnant females, yielding four transgenic pups. The total copy number of the *TH* gene was determined for all pups born to the host mothers using quantitative PCR (data not shown). A pup carrying six total copies of the *TH* gene was assigned as the founder to a “high-copy” line, TH-HI. The TH-HI mice carry two endogenous alleles, as well as four additional copies of *TH* resulting from integration of the transgene (*P* ≤ 0.0001, t-test)(Figure 1A). Commensurate with a threefold increase in gene copy number, *TH* mRNA levels in the midbrain of transgenic mice were three times higher than that of wildtype littermates (measured relative to a commonly-used reference gene, *PGK1*; *P* ≤ 0.0001, t-test) (Figure 1B). TH-HI mice did not differ in size or appearance relative to sex-and age-matched wildtype littermates, nor did they have apparent differences in general health. Mice did not have difficulties breeding or delivering pups, and had an average litter size of 5.2 pups (breeding pairs were retired at 8-12 months of age).

**Figure 1:**
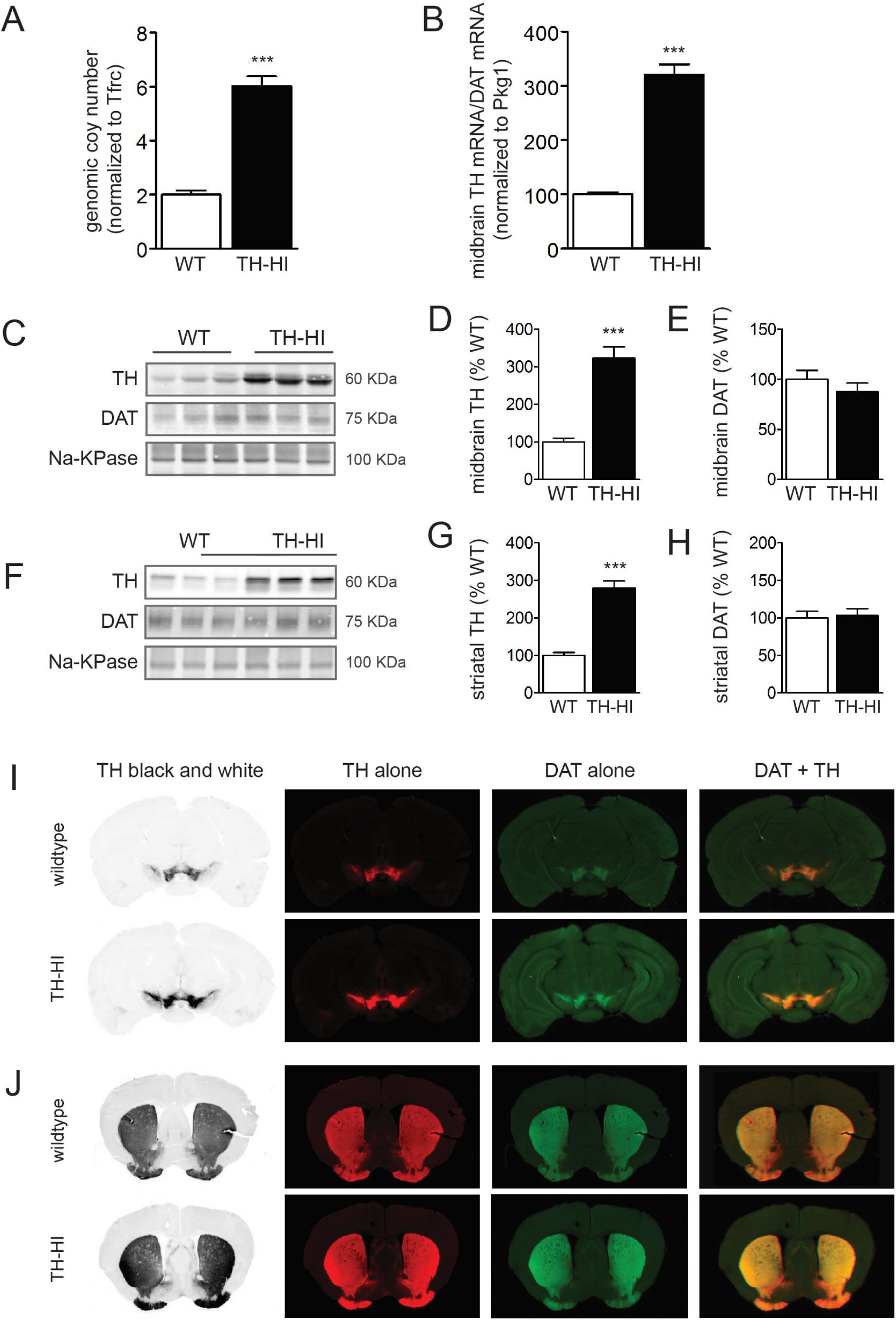
The development of tyrosine hydroxylase overexpressing mice (TH-HI). **(A)** Genomic copy number of tyrosine hydroxylase (TH) in TH-HI mice relative to wildtype (WT) littermates. Copy number is determined by comparison of the cycle threshold to that of the reference gene, *Tfrc*. (n=5 per group)(t(8) = 10.561, *P* ≤ 0.0001). **(B)** *TH* mRNA levels in TH-HI and WT mice, relative to the reference gene *Pkg1*. All values are normalized to dopamine transporter (DAT) mRNA levels in the same sample, which is used as a marker of dopaminergic cells, and shown relative to wildtype levels (WT n=8, TH-HI n=7) (t(13) = 12.41, *P* ≤ 0.0001). **(C)** Midbrain TH and DAT protein expression is shown in adult mice by western blot together with the loading control, Na-KPase (representative samples are shown, n=6 per group). **(D)** Average optical density of midbrain TH protein in TH-HI mice (normalized to Na-KPase), shown relative to wildtype mice (TH, t(10) = 7.158, *P* ≤ 0.0001). **(E)** Average optical density of midbrain DAT protein in TH-HI mice (normalized to Na-KPase), shown relative to wildtype mice (DAT, t(10) = 1.00, *P* = 0.341). **(F)** Striatal TH and DAT protein expression is shown in adult mice by western blot together with the loading control, Na-KPase (representative samples, n=6 per group). **(G)** Average optical density of striatal TH protein in TH-HI mice (normalized to Na-KPase), shown relative to wildtype mice (t(10) = 8.351, *P* ≤ 0.0001). **(H)** Average optical density of striatal DAT protein in TH-HI mice (normalized to Na-KPase), shown relative to wildtype mice (t(10) = 0.257, *P* = 0.803). **(I)** Immunohistochemistry reveals protein expression patterns of TH (red) and DAT (green) in the midbrain. Yellow represents co-localized TH and DAT. **(J)** Protein expression patterns of TH (red) and DAT (green) in the striatum. Yellow represents co-localized TH and DAT. Mean+/−SEM. (*P* < 0.001, ***)

### Total TH Protein Expression Is Increased in Central And Peripheral Sites Of Catecholamine Production

Total TH protein levels were measured to determine if protein levels corresponded to the level of mRNA increase in the transgenic mice. Protein levels were quantified by western blot in juvenile mice (4 weeks of ages) as well as in adult mice (approximately 12 weeks of age; range, 11.5-14 weeks). At both ages, TH-HI mice showed a threefold increase in the optical density of TH protein in the midbrain and striatum, proportional to the observed increase in mRNA (0.0001 ≤ *P* ≤ 0.05 for midbrain and striatum at both ages, t-test) (Figure 1C-D, midbrain at 12 weeks; Figure F-G, striatum at 12 weeks; 4-week, data not shown). No significant change in the level of DAT protein was detected at either time point between wildtype and TH-HI mice (Figure 1E, midbrain at 12 weeks, *P* = 0.32, t-test; Figure 1H, striatum at 12 weeks, *P* = 0.80, t-test; 4-week data not shown). Importantly, TH protein was expressed in similar patterns in the midbrain and striatum of wildtype and transgenic mice, shown using immunohistochemistry (Figure 1J-I). There was no evidence of ectopic expression of TH in the regions that surround the striatum in the forebrain, or the substantia nigra and ventral tegmental area in the mesencephalon. These results are consistent with the advantages of BAC transgenesis, which permits native spatiotemporal protein expression.

To assess protein expression in the noradrenergic system, we first quantified total TH protein in the locus coeruleus, the main source of central noradrenaline. There was an approximate threefold increase in TH optical density, comparable to the increased observed in dopaminergic regions (*P* = 0.00028, t-test) (Figure 2A-B). TH expression patterns in the locus coeruleus of transgenic mice mirrored that of wildtype littermates, again having no detectable ectopic expression in the surrounding regions (Figure 2C).

**Figure 2:**
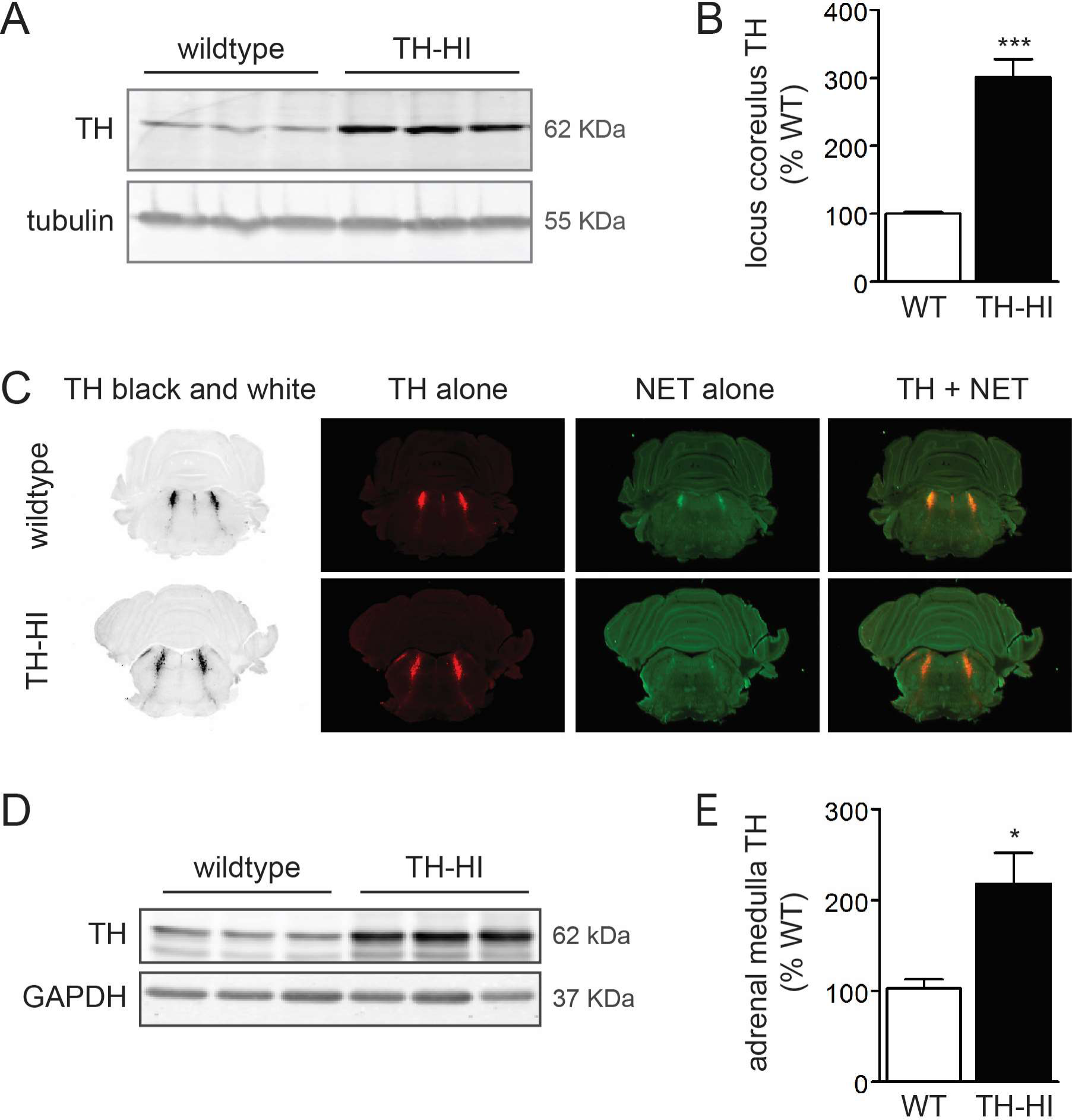
Tyrosine hyroxylase protein expression in central and peripheral noradrenergic regions. **(A)** Protein expression of tyrosine hydroxylase (TH) in the locus coeruleus, with β-tubulin as a loading control (representative samples are shown, WT n=4, TH-HI n=5). **(B)** Optical density of TH in the locus coeruleus (normalized to β-tubulin), shown relative to wildtype levels. (t(7)=6.788, *P* = 0.0003). **(C)** Immunohistochemistry reveals protein expression of TH (red), the noradrenaline transporter (NET, green), and co-labelling (yellow). (Representative samples.) **(D)** Protein expression of TH in the adrenal medulla (representative samples), normalized to GAPDH. **(E)** Optical density of TH in the adrenal medulla (normalized to GAPDH), shown relative to wildtype levels. (WT n=5, TH-HI n=6)(t(9)=2.963, *P* = 0.016) Mean +/− SEM (*P* ≤ 0.05, *; *P* ≤ 0.001, ***).

The expression level of TH was next measured in the periphery. The adrenal medulla is a concentrated site of peripheral TH, secreting noradrenaline and adrenaline in response to input from the autonomic nervous system(de Diego et al., 2008). In transgenic mice, total TH protein in the adrenal medulla was approximately 2.5 times higher than that of wildtype littermates (*P* = 0.016, t-test), demonstrating successful overproduction of TH both in the periphery and the central nervous system (Figure 2D-E).

### Increased Protein Expression Results in Elevated TH-Enzymatic Activity in TH-HI Mice

Activity of TH is modulated by phosphorylation of three main serine (Ser) residues on its amino terminus. Phosphorylation of Ser31 and Ser40 directly affect the activity of TH, while phosphorylation of Ser19 is believed to facilitate enzymatic activity indirectly by encouraging phosphorylation at Ser40 (Haycock, 1993; Lindgren et al., 2000; Dunkley et al., 2004; Daubner et al., 2011). Western blot analysis shows a near-threefold increase in the optical density of phosphorylated TH in the striatum of TH-HI mice at all three relevant residues (Ser40, *P* ≤ 0.0001; Ser31, *P* = 0.010; Ser19, *P* ≤ 0.0001, t-test for all)(Figure 3A-F). This increase in phospho-TH is proportional to the increase in total TH protein seen in transgenic mice (Figure 1C and F). These results indicate that the overexpressed TH is in an active state, and in a conformation less susceptible to feedback inhibition by catechols (Haycock et al., 1982; Haycock, 1987; Haycock and Haycock, 1991; Daubner et al., 2011).

**Figure 3:**
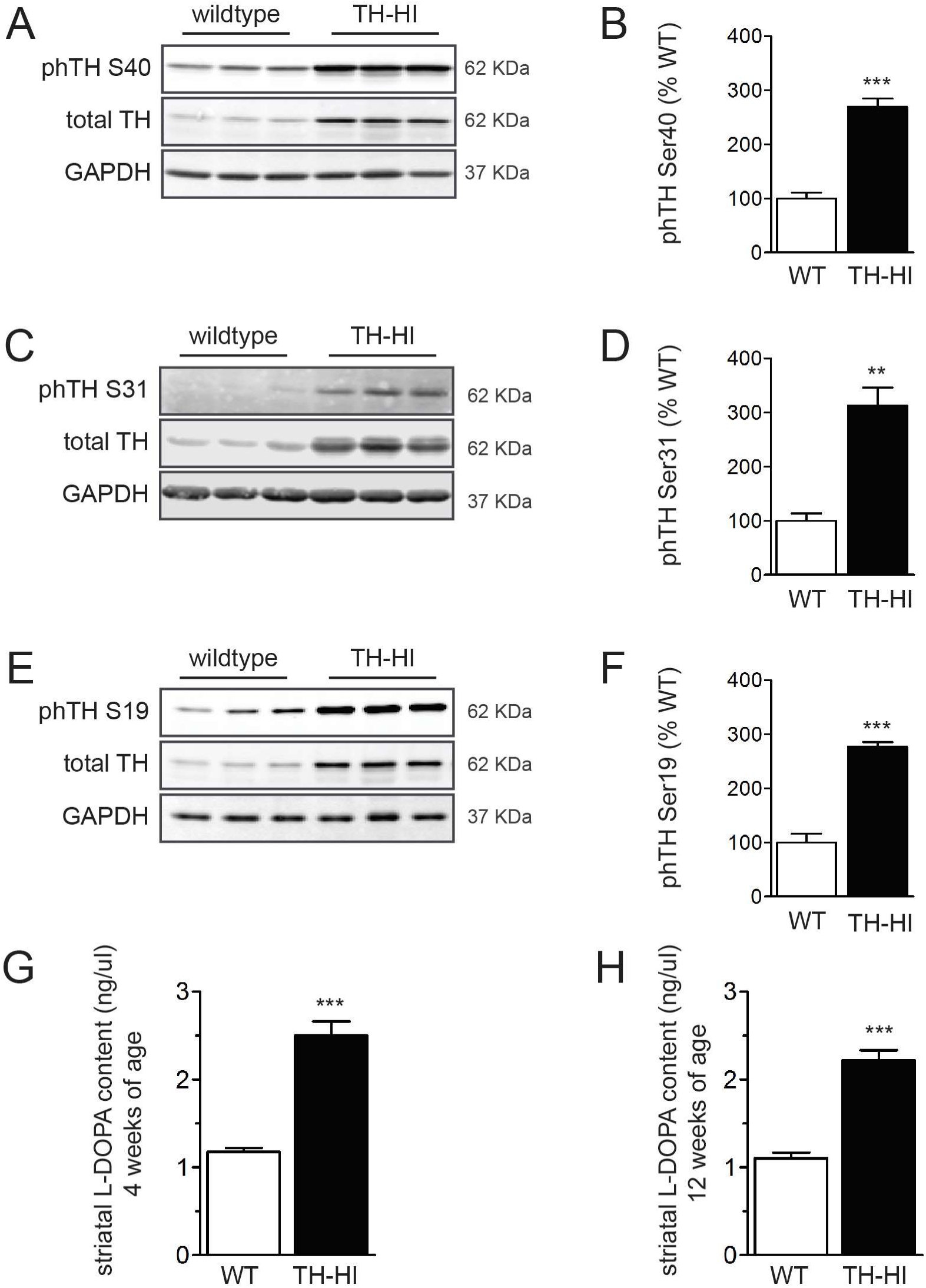
Phosphorylation of TH and L-3,4-dihydroxyphenylalanine (L-DOPA) accumulation in TH-HI mice. Phosphorylation at three main residues affecting activity of tyrosine hydroxylase (TH) was evaluated with western blot. **(A)** Levels of TH protein phosphorylated at serine 40 (phTH S40) (representative samples shown, n=6 per group). **(B)** Optical density of phTH S40 normalized to GAPDH, shown relative to wildtype levels (t(10)=8.942, *P* ≤ 0.0001). **(C)** Levels of TH protein phosphorylated at serine 31 (phTH S31) (representative samples, WT n=4, TH-HI, n=6). **(D)** Optical density of phTH S31 normalized to GAPDH, shown relative to wildtype levels (t(8)=5.010, *P* = 0.0010). **(E)** Levels of TH protein phosphorylated at serine (phTH S19) (representative samples shown, n=6 per group). **(F)** Optical density of phTH S19 normalized to GAPDH, shown relative to wildtype levels (t(10)=9.687, P ≤ 0.0001) **(G)** L-DOPA accumulation following injections of NSD-1015 at 4-weeks of age, detected using high-performance liquid chromatography (WT n=9, TH-HI n=8; P ≤ 0.0001). **(H)** L-DOPA accumulation following NSD-1015 injections at 10-16 weeks of age (WT n=7, TH-HI n=7; *P* ≤ 0.0001). Mean +/− SEM (*P* ≤ 0.01, **; *P* ≤ 0.001, ***).

We next sought to confirm that the overexpressed TH protein is functional by measuring striatal L-DOPA levels, which is the direct product of TH. In both 4 week-old and adult TH-HI mice, L-DOPA accumulation was 200% that of wildtype levels following injections of NSD-1015 (*P* ≤ 0.0001 for both ages, t-test) (Figure 3G-H). This twofold increase in L-DOPA production directly demonstrates that the overexpressed TH protein is functionally active *in vivo*.

### Dopamine Turnover Is Increased in TH-HI Mice

As the rate of catecholamine synthesis was shown to be increased by the accumulation of L-DOPA, we next measured the tissue content of dopamine and its metabolites in the striatum of both juvenile and adult mice. Striatal tissue content of dopamine was increased by approximately 23% in 4 week-old mice TH-HI mice compared to wildtype littermates (Figure 4A) (*P* = 0.018, two-way ANOVA); however, there was no detectable change in dopamine tissue content levels in adult mice aged 10-20 weeks (Figure 4A). In contrast, the total tissue content of DOPAC and HVA was significantly higher in both 4 weeks *and* adult mice (Figure 4B). Indeed, striatal content of DOPAC measured in TH-HI mice was 81% over wildtype levels in 4 week-old mice and 52% in adult mice, while the content of HVA increased 63% and 50% in juvenile and adult mice respectively (*P* < 0.001 for all, two-way ANOVA) (Figure 4B, D). The ratios of metabolites to dopamine were also significantly increased over wildtype levels at both ages (Figure 4C and E). In juvenile TH-HI mice, the ratio of DOPAC-to-dopamine increased 51% and the HVA-to-dopamine ratio increased 36% over wildtype levels. These ratios rose to 58% and 59% (respectively) over wildtype levels in adult mice (*P* < 0.001 for all, two-way ANOVA) (Figure 4C,E). Importantly, a high metabolite to dopamine ratio is representative of increased dopamine turnover, previously linked to oxidative stress in dopaminergic striatal terminals as a result of ROS that are produced during the deanimation of dopamine (Spina and Cohen, 1989).

**Figure 4:**
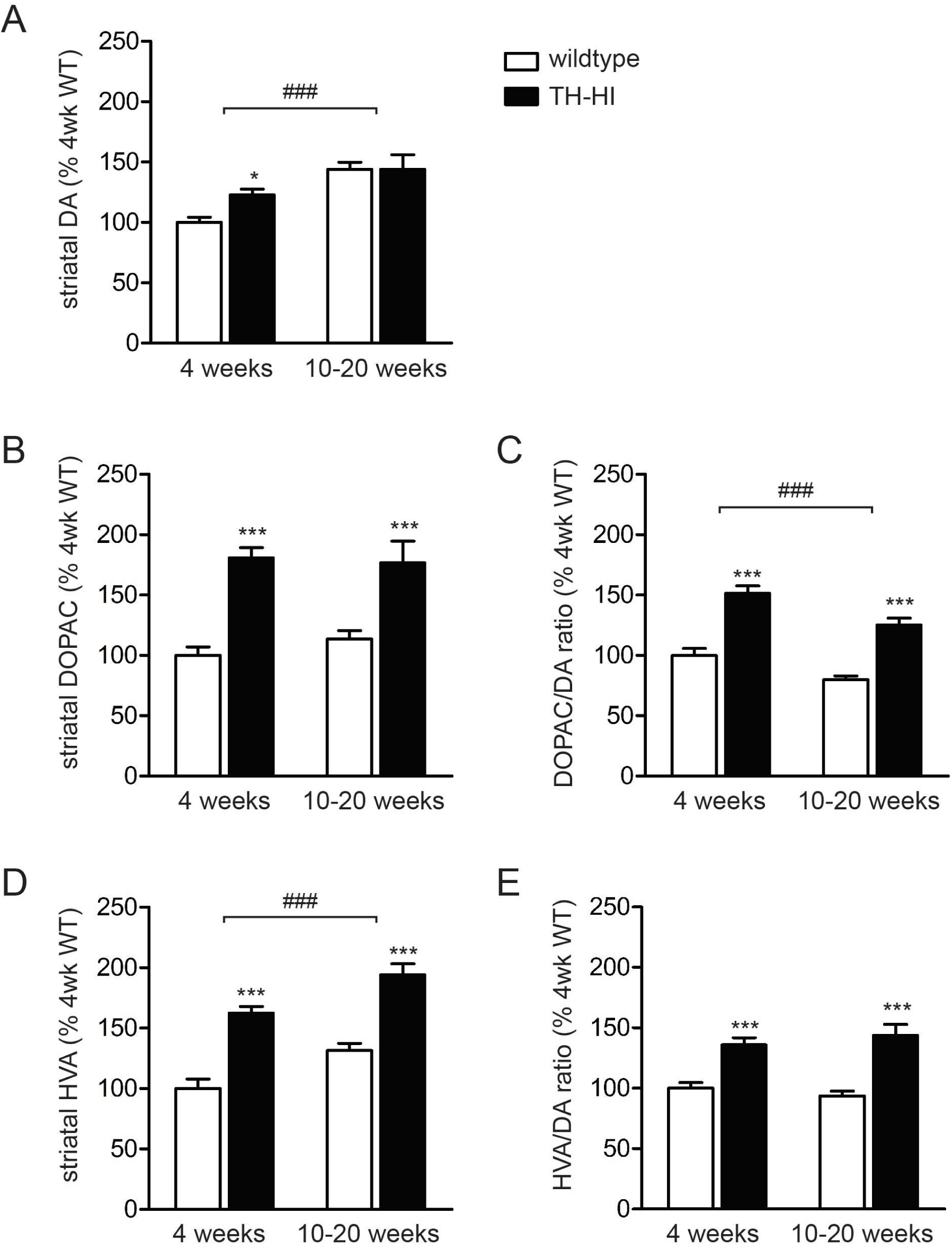
Striatal tissue content of dopamine and its metabolites. In juvenile and adult mice, tissue content of **(*A*)** dopamine (DA) (two-way ANOVA: effect of age, F1,64 = 23.63, *P* ≤ 0.0001); **(*B*)** 3,4-dihydroxyphenylacetic acid (DOPAC) (two-way ANOVA, effect of genotype: F_1,64_ = 53.28, *P* ≤ 0.0001) and **(*D*)** homovanillic acid (HVA) (two-way ANOVA: effect of age, F1,64 = 19.02, *P* ≤ 0.0001; effect of genotype, F1,64 = 74.36, *P* ≤ 0.0001). The ratios of metabolite-to-dopamine were also determined: **(*C*)** DOPAC to DA (two-way ANOVA: effect of age, F1,64 = 18.76, *P* ≤ 0.0001; effect of genotype: F_1,64_ = 82.22, *P* ≤ 0.0001) and **(*E*)** HVA to DA (two-way ANOVA, effect of genotype: F1,64 = 55.37, *P* ≤ 0.0001). (4 weeks: WT n=20, TH-HI n=15; 10-20 weeks: WT n=20, TH-HI n=13) (variation due to age denoted “#”:*P* < 0.001, ###). Mean+/−SEM, normalized to 4-week old mice WT levels for each analyte (Bonferroni post-hoc test: *P* ≤ 0.05, *; *P* ≤ 0.001, ***).

### TH-HI Mice Have a Potentiated Response to Amphetamine That Is Blocked by NSD-1015

To evaluate behavioural consequences of increased TH activity in transgenic mice, we challenged mice with moderate and high-doses of amphetamine (2.0 mg/kg and 3.0 mg/kg, respectively). Amphetamine treatment results in massive efflux of monoamines through their membrane transporters due to a reversal in the concentration gradient, resulting in the stimulation of dopamine-and noradrenaline-controlled functions (Sulzer et al., 1995; Jones et al., 1998a; Salahpour et al., 2008; Heal et al., 2013). Because of its potent effect on the dopaminergic system, amphetamine is often used to test the integrity of the nigrostriatal pathway by using locomotor activity as an outcome measure (Giros et al., 1996). At baseline, the locomotor behaviour of transgenic mice was not significantly different from their wildtype littermates at either of the ages (Figure 5A). However, adult TH-HI mice had a greatly potentiated locomotor response to amphetamine, more than doubling total distance travelled in the 90 minutes following a 2.0 or 3.0 mg/kg dose (Figure 5B and C). In addition, TH-HI mice showed increased stereotypic behaviour in response to the drug, increasing their stereotypic counts twofold following the administration 2.0 or 3.0 mg/kg of amphetamine (Figure 5D). The response of adult mice matches that of four-week old TH-HI mice: following a 3.0 mg/kg intraperitoneal dose of amphetamine, 4-week old mice cover three times the distance of wildtype littermates (18,620 +/− 1384 cm/min in TH-HI compared to 5840 +/− 1027 cm/90min in WT; n=6 for both, *P* ≤ 0.0001) and have a two-fold increase in stereotypic behaviour (11,660 +/− 1163 counts compared to 5690 +/− 745 counts/90 minutes in WT mice; n=6 for both, *P* = 0.0015) (results not shown).

**Figure 5:**
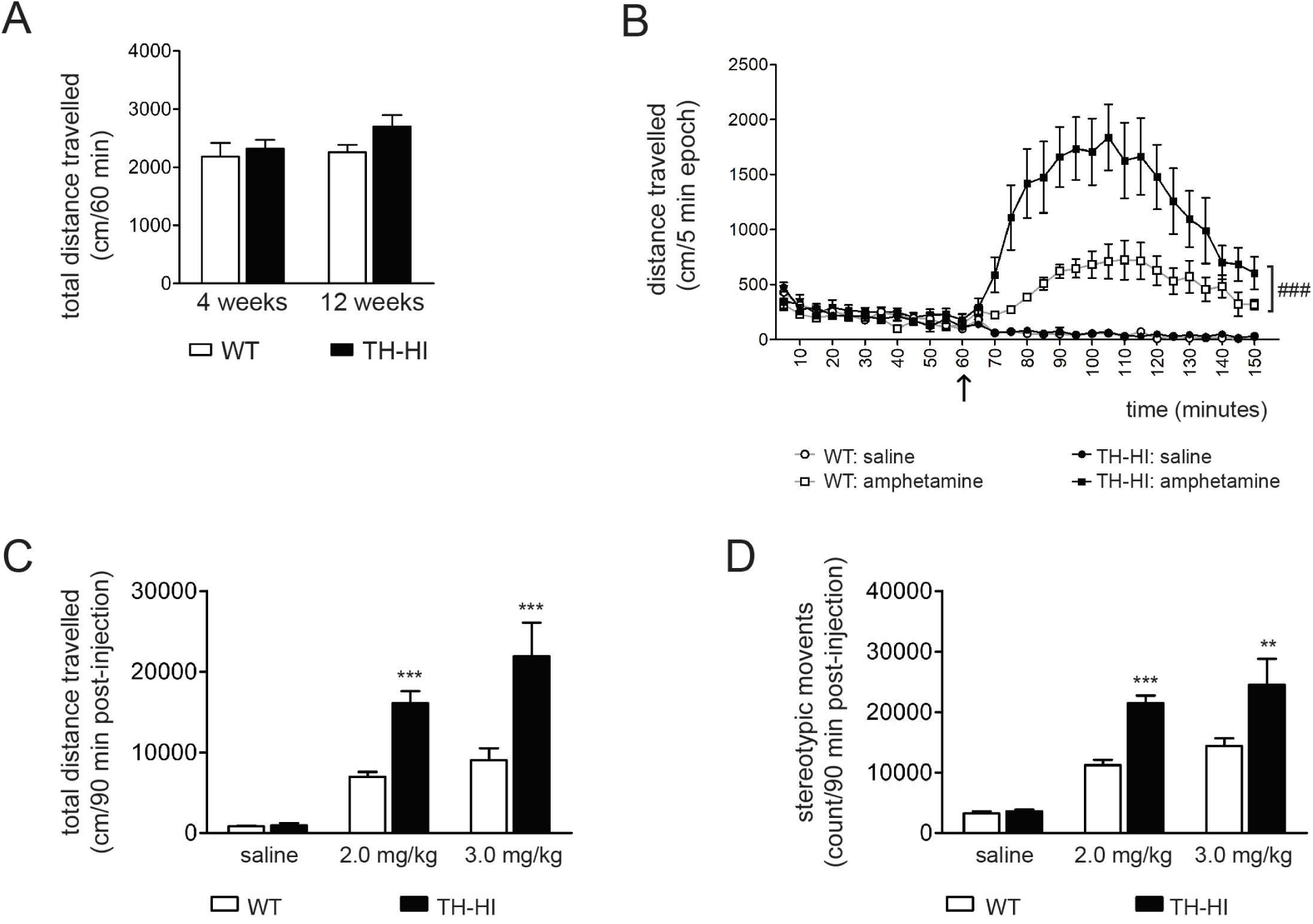
Locomotor activity at baseline and the response to amphetamine. **(A)** Total distance (cm) travelled by wildtype (WT) and transgenic (TH-HI) mice in 60 minutes, at baseline. (Repeated-measures two-way ANOVA: effect of time, F29,551 = 64.79, *P* ≤ 0.0001; effect of genotype, F_1,551_ = 36.14, *P* ≤ 0.0001; genotype x time interaction: F_29,551_ = 17.85, *P* ≤ 0.0001) (4 weeks: n=10 for both groups; 12 weeks: WT n=33, TH-HI n=28) **(B)** Total distance travelled by adult mice in 5 minute epochs, before and after injection with saline or 3.0 mg/kg amphetamine (black solid-lined arrow denotes the time of injection). (Repeated measures two-way ANOVA for saline: effect of time, F29,290 = 34.66, *P* ≤ 0.0001; effect of genotype, F1,290 = 0.01, P = 0.931; genotype x drug interaction, F29,290 = 0.78, *P* = 0.780) (Repeated measures two-way ANOVA for 3.0 mg/kg amphetamine: effect of time, F_29,290_ = 20.33, *P* ≤ 0.0001; effect of genotype, F1,290 = 9.83, P = 0.011; genotype x drug interaction, F29,290 = 5.49, P ≤ 0.0001) (Genotype effect in the amphetamine group shown as ###, *P* < 0.0001) **(C)** Total distance travelled in 90 minutes after injection of saline (Two-way ANOVA: effect of drug, F_2,39_ = 35.36, *P* ≤ 0.0001; effect of genotype, F1,39 = 28.45, *P* ≤ 0.0001; genotype *x* drug interaction, F2,39 = 6.625, *P* = 0.0033) **(D)** Total stereotypic counts in 90 minutes after injection of saline, 2.0 mg/kg, or 3.0 mg/kg amphetamine. (two-way ANOVA: effect of drug, F_2,39_ = 43.40, *P* ≤ 0.0001; effect of genotype, F1,39 = 23.73, *P* ≤ 0.0001; genotype x drug interaction, F2,39 = 5.09, *P* = 0.011) (For B, C and D, saline: WT n=6, TH-HI n=6; 2.0 mg/kg: WT n=13, TH-HI n=8; 3 mg/kg: WT n=6, TH-HI n=6.) Mean +/− SEM (Bonferroni post-hoc: *P* ≤ 0.05, *, *P* ≤ 0.01, **; *P* ≤ 0.001, ***).

### VMAT2 Levels Are Unchanged in TH-HI Transgenic Mice

We next sought to determine if the expression of VMAT2 — the principle protein responsible for packaging monoamines into vesicles — was upregulated in TH-HI mice to accommodate increased TH synthetic activity. Importantly, a change in the expression of VMAT2 can reflect one strategy by which the cell might mitigate damage induced by increased production of monoamines. Figure 6 shows that there was no difference in VMAT2 protein levels between TH-HI mice and wildtype littermates at 4 weeks and 12 weeks of age (Figure 6A-B and Figure 6 C-D, respectively).

**Figure 6:**
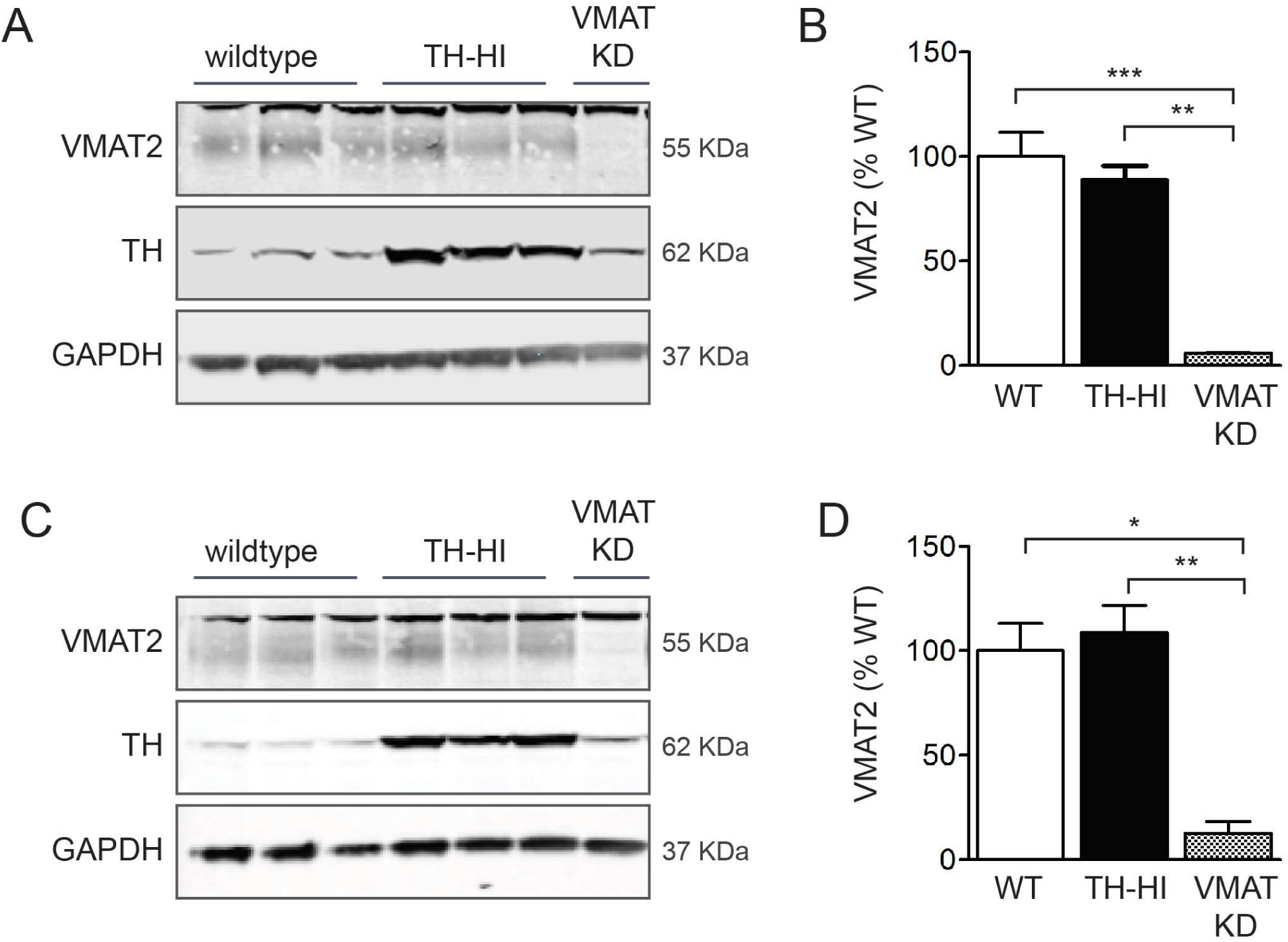
Striatal protein levels of the vesicular monoamine transporter (VMAT2) in juvenile and adult mice. **(A)** Representative samples showing VMAT2 at 4 weeks of age and **(B)** optical density of VMAT2, normalized to GAPDH (WT n=6; TH-HI n=6; VMAT-kd, n=2) (F_2,13_ = 14.19, *P* = 0.0009). (***C***) Representative samples showing VMAT2 at 12 weeks of age and (***D***) optical density of VMAT2, normalized to GAPDH (WT n=5; TH-HI n=6; VMAT-kd, n=2) (F2,12 = 8.43, *P* = 0.0072). (one-way ANOVA, Bonferroni post-hoc test: *P* ≤ 0.05, *; *P* ≤ 0.01,**; *P* ≤ 0.001,***) Mean +/− SEM.

### TH-HI Have Markers of Oxidative Stress

We have shown that in TH-HI mice, there is a significant and sustained increase in L-DOPA production as well as an increase in dopamine turnover, despite no measurable increase in tissue content of dopamine in adult mice. Importantly, VMAT2 levels were unchanged in TH-HI mice. These results suggest that additional dopamine may indeed be produced in adult mice but not immediately sequestered, giving it increased opportunity to be degraded to DOPAC and HVA. Because hydrogen peroxide (H2O2) is produced by monoamine oxidase during the metabolism of dopamine (Maker et al., 1981; Spina and Cohen, 1989; Halliwell, 1992), an elevated presence of metabolites could indicate accelerated ROS production in TH-HI mice. To evaluate the presence of oxidative stress in adult TH-HI mice, we first examine levels of glutathione, which neutralizes H2O2 by converting it to H2O (Spina and Cohen, 1989). Figure 7A-B shows that levels of glutathione were significantly reduced in the striatum of adult TH-HI mice but not in cortical tissue, indicating that there is an elevated presence of H2O2 in the striatum, where increased TH activity is observed.

**Figure 7:**
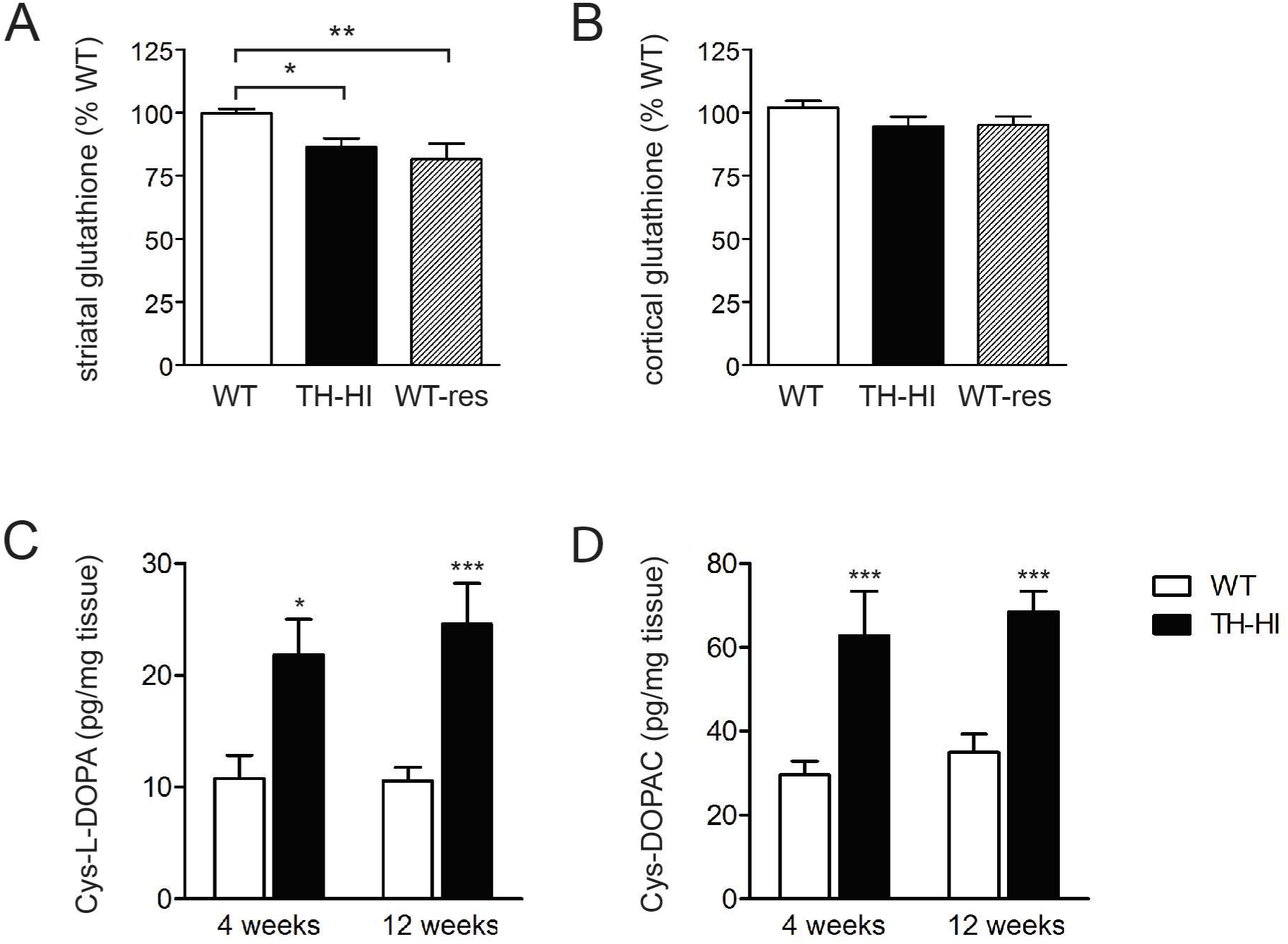
Markers of oxidative stress elevated in adult TH-HI mice. (***A***) Total glutathione levels (GSH) were reduced in striatal tissue of adult transgenic mice (TH-HI) as compared to wildtype (WT) (12-15 weeks of age). No significant difference in striatal GSH levels was detected between transgenic mice and WT mice that had been treated with reserpine two hours prior to dissections (WTres). (WT n=14; TH-HI n=14; WTres n=7) (all data normalized to WT) (one-way ANOVA: F 2, 32 = 7.709, *P* = 0.0019) (***B***) No change in GSH levels was detected in cortical samples. (WT n=14; TH-HI n=14; WTres n=7). (All data normalized to WT.) Mean +/−SEM. (**C**) Cysteinylated L-DOPA levels (Cys-L-DOPA) in the striatum of TH-HI and wildtype mice at 4 weeks and 12 weeks of age (4 weeks: WT n=10, TH-HI n=9; 12 weeks: WT n=14, TH-HI n=12) (two-way ANOVA: F 1, 41 = 22.87, *P* ≤ 0.0001) (**D**) Cysteinylated DOPAC levels (Cys-DOPAC) in the striatum of TH-HI and wildtype mice at 4 weeks and 12 weeks of age (4 weeks: WT n=10, TH-HI n=7; 12 weeks: WT n=12, TH-HI n=13) (two-way ANOVA: F 1, 38 = 35.29, *P* ≤ 0.0001). (Bonferrori post-hoc, all pairwise comparisons: *P* ≤ 0.05, *; *P* ≤ 0.01, **; *P* ≤ 0.001, ***) Mean +/− SEM.

In addition to undergoing metabolism in the cytosol, unsequestered dopamine and DOPAC are susceptible to oxidation, forming ROS and *o*-quinones (Graham, 1978; Hastings et al., 1996; Chen et al., 2008). These oxidation products can then react with reduced sulfhydryl groups such as free cysteine or glutathione, or covalently bind to cysteine groups on nucleophilic proteins (often at the active site), which can impair their function and have deleterious effects on the cell (Hastings and Zigmond, 1994; Hastings et al., 1996). Therefore, the presence of cysteinyl catechols is used as a measure of the oxidation of free cytosolic dopamine and its metabolites in the dopaminergic regions of the brain (Hastings and Zigmond, 1994; Hastings et al., 1996; Caudle et al., 2007; Chen et al., 2008). We measured cysteinyl-DOPA and cysteinyl-DOPAC in juvenile and adult mice to assess whether TH-overexpression resulted in an increased presence of cysteinylated catechols. Corroborating results that show TH-HI mice have reduced gluthathione levels, we found an increased presence of both cysteinyl-DOPA and cysteinyl-DOPAC in TH-HI animals (Figure 7C and D). Cysteinyl-L-DOPA and cysteinyl-DOPAC levels were increased approximately 200% in 4-week old TH-HI compared to age-matched controls. Adult TH-HI mice had similarly elevated cysteinyl conjugates compared to wildtype littermates, again showing a twofold increase in cysteinyl-L-DOPA and cysteinyl-DOPAC levels. These results demonstrate that markers of oxidative stress are present in the striatum of mice with elevated TH activity.

## Discussion

While previous efforts to develop TH-overexpressing mice have been made (Kaneda et al., 1991), the mice presented in this paper are the first that successfully over-produce functional TH in endogenous catecholaminergic regions and this study is the first to report the consequences of increased TH expression in transgenic mice. These transgenic mice possess six total copies of the murine *TH* gene with a commensurate increase in mRNA and phosphorylated TH, indicative of a proportional increase in active-state enzyme (Figures 1, 2, and 3A-F). Here, we show that directly increasing TH activity is sufficient to produce markers of oxidative stress.

Past studies have demonstrated *in vitro* that TH in its active state can contribute to the formation of ROS as can the related enzyme phenylalanine hydroxylase (Adams et al., 1997). As with many enzymes, phosphorylation plays an important role in regulating its activity levels (Haavik et al., 1989; Dunkley et al., 2004). Indeed, the phosphorylation of TH induces a conformational change releasing it from end-product inhibition, in which catechols are bound to iron in its ferric (Fe^3+^) state following a hydroxylation reaction; the unbinding of catechols allows the iron molecule to be reduced to its ferrous (Fe^2+^) form, returning TH to an active state and freeing the iron molecule for further interactions (Dunkley et al., 1996; Haavik, 1997; Haavik et al., 1997). In this way, an increased level of phosphorylated TH directly influences the rate of catecholamine synthesis, as we observed in TH-HI mice (Figure 3). However, *in vitro* studies have also linked oxidative stress with the TH biosynthetic system, having demonstrated that ROS such as superoxide and H2O2 can be created during catalytic turnover (Adams et al., 1997; Haavik et al., 1997). In addition, results from *in vitro* studies suggest that the Fe^2+^ molecules bound in TH’s active-site has the potential to further participate in a Fenton-like reaction, interacting with H_2_O_2_ to form highly reactive hydroxyl radicals (Adams et al., 1997; Haavik, 1997; Haavik et al., 1997). Therefore, that TH-HI mice have a threefold increased levels of phosphorylated TH is a meaningful observation. Not only does it signify that the additional TH produced in transgenic mice is active and capable of increased catecholamine synthesis, but it may also indicate a greater potential for TH to participate in the formation of reactive species.

Juvenile and adult TH-HI mice showed increased rates of dopamine synthesis compared to age-matched littermates, reflected by increased L-DOPA accumulation following NSD-1015 administration (Figure 3G-H). While HPLC detected an elevated level of striatal tissue content of dopamine at 4 weeks only, TH-HI mice have a sustained increase in striatal content of DOPAC (Figure 4B) and HVA (Figure 4D) at both ages. The increased presence of striatal dopamine metabolites in the absence of a corresponding increase in dopamine tissue content in adult animals suggests that the additional dopamine produced in transgenic mice may disproportionally reside in the cytosol, where it is quickly metabolised (Spina and Cohen, 1989). Under normal circumstances, the majority of the catechoamine pool is stored in vesicles (Fon et al., 1997). Figure 6 shows that VMAT2 levels in TH-HI mice do not differ from wildtype littermates, which may be an indication that excess dopamine produced in transgenic mice has a greater potential to be metabolised more quickly than it can be sequestered. Together, these findings are consistent with studies showing that VMAT2 knockout (VMAT2-/-) mice have normal levels of DOPAC and HVA in the striatum despite having just 1.5% of wildtype dopamine levels, which was attributed to rapid metabolism of transmitter that cannot be stored in vesicles (Fon et al., 1997).

TH-HI mice show a potentiated locomotor response to amphetamine, showing a high magnitude of response at both 4 and 12 weeks of age. Stored dopamine is released from vesicles upon exposure to amphetamine, joining freshly synthesized and recycled dopamine in the cytosol. The increased concentration of dopamine causes a reversal in the directionality of DAT, allowing dopamine to be transported down its concentration gradient into the extracellular space (Sulzer et al., 1995). However, using cultured midbrain neurons, Fon et al. (1997) showed that the amphetamine response is not dependent on vesicular stores of dopamine, and can results from efflux of non-vesicular (cytosolic) dopamine through plasma membrane transporters. Previous studies have also shown amphetamine-induced flux reversal across the plasma membrane increases TH activity by eliminating feedback inhibition by cytosolic catechols, particularly when vesicular stores are depleted (Spector et al., 1967; Zigmond et al., 1989). Using VMAT2-/-mice, Fon et al. (1997) further demonstrated that the *de novo* synthesis of dopamine can play a large role in the behavioural response under abnormal conditions, reporting that amphetamine administration in neonatal VMAT2-/- mice promoted locomotor activity and sustained complex behaviours, even in the absence of vesicular stores of dopamine. Here, we show that TH-HI have a twofold increase in the locomotor response to amphetamine compared to wildtype mice, with an increase in both the total distance travelled and the stereotypic response to the stimulant at both ages. That a twofold increase in striatal dopamine metabolites was observed in TH-HI mice suggests that at baseline conditions, excess dopamine is produced and quickly degraded; however, through a rapid amphetamine-induced release into the extracellular space, its presence is revealed.

Accelerated dopamine turnover in the cytosol has been shown to alter the redox status of presynaptic dopamine neurons, and has long been identified as a source oxidative stress in the nigrostriatal pathway (Spina and Cohen, 1989). In presynaptic terminals, unsequestered dopamine undergoes degradation by monoamine oxidase (a mitochondrial enzyme) to form metabolites as well as H2O2 (Graham, 1978; Stokes et al., 1999; Hermida-Ameijeiras et al., 2004). While the degradation of dopamine serves to protect a cell from intracellular damage resulting from its auto-oxidation, accelerated metabolism can also become a source of oxidative stress. Normally, small amounts of oxygen species produced during catalytic or metabolic reactions is scavenged by reduced-form glutathione and superoxide dismutase (Spina and Cohen, 1989; Dunnett and Bjorklund, 1999). In this study, mice overexpressing active TH show increased levels of dopamine turnover (Figure 4C and E) that is accompanied by a decrease in the reduced-form of glutathione in the striatum (Figure 7A), indicating the presence of oxidative stress (Spina and Cohen, 1989). Not only are these findings consistent with other animal models of dopamine toxicity (Hastings et al., 1996; Caudle et al., 2007; Chen et al., 2008), but they are congruent with post-mortem analyses of the brains of Parkinson’s disease patients, where increased oxidative damage and mitochondrial complex I deficiencies have been reported in substantia nigra and frontal cortex (Schapira, 1993; Keeney et al., 2006; Henchcliffe and Beal, 2008; Parker et al., 2008; Moon et al., 2013). Additionally, decreased level of reduced-form of glutathione has also been reported in the substantia nigra of Parkinson’s disease patients compared to age-matched controls (Perry and Yong, 1986; Sian et al., 1994).

Together with changes in reduced glutathione, we also observed profound increases in cysteinyl-DOPAC and cysteinyl-DOPA in juvenile and adult TH-HI mice, demonstrating accelerated oxidation. The relative amount of free cysteinyl catechols has been shown to increase under conditions that promote oxidative stress, including high levels of cytosolic dopamine, reserpine treatment, and even normal ageing (Fornstedt et al., 1989; Fornstedt and Carlsson, 1989; Fornstedt et al., 1990; Hastings et al., 1996; Chen et al., 2008). Our results show an elevated presence of oxidized L-DOPA and dopamine metabolites in mice that have increased activity of TH, consistent with models of intracellular dopamine mishandling that have previously linked cytosolic dopamine to oxidative stress (Caudle et al., 2007; Masoud et al., 2015).

In this paper, we have shown that increased levels of active-form TH leads to an accelerated rate of dopamine turnover and oxidative stress in dopamine-rich brain regions. This could represent a source of oxidative stress specific to catecholamine cells in the early stages of pathogenesis of Parkinson’s disease. Further linking our studies to Parkinson’s disease is the observation that in alpha-synuclein knockout mice, lentiviral transduction of human alpha-synuclein results in increased TH phosphorylation in cells expressing alpha-synclein aggregates (Alerte et al., 2008). Moreover, studies in *Drosophilia, C elegans*, and in stably transfected MN9D cells, have shown that expression of mutated alpha-synuclein results in increased TH activity, dopamine production, oxidative stress and neurodegeneration (Perez et al., 2002; Park et al., 2007; Locke et al., 2008). Early pathological changes in Parkinson’s disease, and the development of alpha-synuclein aggregates, may therefore provide an environment that facilitates further oxidative stress and cellular damage arising from increased activity of tyrosine hydroxylase. Our findings support further study of TH as a potential contributor to catecholamine dysregulation and oxidative stress and, perhaps a target for therapeutic intervention.

## Acknowledgments

We gratefully acknowledge Wendy Horsfall for animal husbandry and mouse injections, as well as Lauren Tessier for technical assistance. This research was supported by Canadian Institutes of Health Research operating grants 210296 to AS and 258294 to AJR. We also thank the Government of Ontario for their support through Ontario Graduate Scholarships and the Queen Elizabeth II/Grace Lumsden/Margaret Nicholds Graduate Scholarship in Science and Technology.

